# Anti-microbial immunity is impaired in COPD patients with frequent exacerbations

**DOI:** 10.1101/632372

**Authors:** Aran Singanayagam, Su-Ling Loo, Maria Calderazzo, Lydia J Finney, Maria-Belen Trujillo Torralbo, Eteri Bakhsoliani, Jason Girkin, Punnam Veerati, Prabuddha S Pathinayake, Kristy S Nichol, Andrew Reid, Joseph Foottit, Sebastian L Johnston, Nathan W Bartlett, Patrick Mallia

**Author notes:** These authors contributed equally to this work (Sebastian L Johnston, Nathan W Bartlett, Patrick Mallia). **Corresponding author**: Dr Aran Singanayagam. National Heart and Lung Institute. Imperial College London.

## Abstract

**Background:** Patients with frequent exacerbations represent a chronic obstructive pulmonary disease (COPD) sub-group requiring better treatment options. The aim of this study was to determine the innate immune mechanisms that underlie susceptibility to frequent exacerbations in COPD.

**Methods:** We measured sputum expression of immune mediators and bacterial loads in samples from patients with COPD at stable state and during virus-associated exacerbations. *Ex vivo* immune responses to rhinovirus infection in differentiated bronchial epithelial cells (BECs) sampled from patients with COPD were additionally evaluated. Patients were stratified as frequent exacerbators (≥2 exacerbations in the preceding year) or infrequent exacerbators (<2 exacerbations in the preceding year) with comparisons made between these groups.

**Results:** Frequent exacerbators had reduced sputum cell mRNA expression of the anti-viral immune mediators type I and III interferons and reduced interferon-stimulated gene (ISG) expression when clinically stable and during virus-associated exacerbation. RV-induction of interferon and ISGs *ex vivo* was also impaired in differentiated BECs from frequent exacerbators. Frequent exacerbators also had reduced sputum levels of the anti-microbial peptide mannose-binding lectin (MBL)-2 with an associated increase in sputum bacterial loads at 2 weeks following virus-associated exacerbation onset. MBL-2 levels correlated negatively with bacterial loads during exacerbation.

**Conclusion:** These data implicate deficient airway innate immunity in the increased propensity to exacerbations observed in some patients with COPD. Therapeutic approaches to boost innate antimicrobial immunity in the lung could be a viable strategy for prevention/treatment of frequent exacerbations.

## INTRODUCTION

Chronic obstructive pulmonary disease (COPD) is an inflammatory airway disorder in which acute exacerbations represent a major complication. Acute exacerbations are a substantial cause of morbidity and mortality^1–3^ and preventing exacerbations remains a major unmet need. There has been increasing interest in a recently identified sub-group of patients with COPD who are at risk of exacerbations, defined as the ‘frequent exacerbator phenotype’.^4^ Patients with frequent exacerbations have worse clinical outcomes including increased morbidity^1^, accelerated lung function decline^2^ and greater mortality^5^, suggesting that this group requires special consideration. To date, the underlying mechanisms that predispose such patients to more frequent exacerbations have not been elucidated. It is plausible that abnormalities in host immunity could underlie an increased propensity to exacerbations.

Viruses are a major aetiological trigger for exacerbations^6,7^ and data exists to suggest that COPD may be associated with deficient anti-viral immunity^8,9^. However, not all studies have shown this abnormality^10,11^, suggesting that it may vary according to disease phenotype. Frequent exacerbators could represent one sub-group in whom defective anti-viral immunity is more prominent. Both experimental and naturally-occurring exacerbation studies also confirm that an initial virus infection can precipitate secondary bacterial infection in COPD^12,13^ with neutrophil elastase-induced cleavage of anti-microbial peptides believed to be important mechanistically^12^. Although bacterial colonisation at stable state has been shown to be associated with increased exacerbation frequency^14^, antimicrobial peptide responses and propensity to develop secondary bacterial infection during virus-infections in frequent versus infrequent exacerbators has not previously been studied.

We hypothesised that patients with COPD who have a history of frequent exacerbations would have impaired anti-viral and/or anti-bacterial immunity compared to infrequent exacerbators. Using analysis of sputum samples from a cohort of patients monitored prospectively in the community, in combination with *in vitro* experiments in BECs sampled from patients with COPD and differentiated at the air-liquid interface (ALI), we show that lung innate anti-microbial immunity is impaired in frequent exacerbators. These mechanisms may underlie the increased propensity to exacerbation observed in some patients.

## METHODS

### The St Mary’s Hospital naturally-occurring COPD exacerbation cohort

A cohort of 40 COPD subjects was recruited to a longitudinal study carried out at St Mary’s Hospital, London UK between June 2011 to December 2013 investigating the pathogenesis of naturally-occurring COPD exacerbations. The study protocol was approved by the East London Research Ethics Committee (study number 11/LO/0229) and all subjects gave informed written consent. The subjects all had a clinical diagnosis of COPD that was confirmed with spirometry. All subjects had an initial visit at baseline when clinically stable for clinical assessment, spirometry (forced expiratory volume in 1 second (FEV_1_), forced vital capacity (FVC) and peak expiratory flow (PEF)) and clinical sample collection including spontaneous or induced sputum, taken and processed as previously described^8,15^. At the baseline visit, all subjects were asked about the number of exacerbations experienced in the previous year and were grouped into two groups – frequent exacerbators (≥2 exacerbations in preceding year) and infrequent exacerbators (0 – 1 exacerbations in the previous year), as previously described^16^. Subjects then had repeat visits at three monthly intervals when clinically stable and were followed up for a minimum of 6 months. Subjects reported to the study team when they developed symptoms of an upper respiratory tract infection or an increase in any of the symptoms of dyspnoea, cough, sputum volume or purulence. Exacerbation was defined using the East London cohort criteria.^2^ Subjects were seen within 48 hours of onset of their symptoms for collection of samples and repeat visits was carried out a two weeks after the initial exacerbation visit. Samples from this cohort have been used in a previous study investigating the role of airway glucose in COPD^17^

### Experimental challenge studies

For baseline analyses of sputum soluble mediators, in addition to samples taken from the naturally occurring exacerbation study, we also included baseline (pre-infection) samples from 14 COPD subjects recruited to experimental challenge COPD studies^8,18^. All subjects in these studies gave informed written consent and the protocol was approved by St Mary’s NHS Trust Research Ethics Committee (study numbers 00/BA/459E and 07/H0712/138). These subjects were characterised as frequent or infrequent exacerbators using the same definition as described above. Sputum samples were processed using exactly the same methodology as described in the naturally occurring exacerbation study.

### RV infection of air-liquid interface differentiated bronchial epithelial cells from patients with COPD

Primary BECs obtained bronchoscopically from patients with COPD (GOLD Stage II or III) were grown until confluent and differentiated at air liquid-interface (ALI), as previously described^19–21^. All subjects included gave informed written consent and the study protocol was approved by the Hunter New England Human Research Ethics Committee (05/08/10/3.09). Primary cells were grown in complete bronchial epithelial cell growth medium (BEGM)(Lonza) with growth factor supplements in submerged monolayer culture and then seeded at 2 x 10^5^ cells/mL in transwells until confluent. RV-A1 infection (MOI 0.1) was applied to the apical surface for 2 hours in 250μl bronchial epithelial basal medium (BEBM) minimal at 35°C. Following this, infection media was replaced with 500μl fresh BEBM minimal for the remainder of the experiment. Samples were collected at 72 hours post-infection with apical media removed and stored for protein expression analyses. Type III IFN proteins were measured using a custom-designed LEGENDplex kit (BioLegend). IFN-β was measured using the Verikine ELISA (PBL). Half of the transwell membrane was also carefully cut from the inserts and collected into RLT buffer (Qiagen) containing 1% 2-mercaptoethanol for downstream molecular analyses by RT-qPCR. The remaining transwell membrane was fixed in 10% neutral-buffered formalin for 24 hours for histological cross-sections to confirm differentiation.

### Measurement of sputum proteins

Protein levels in cell-free sputum samples of CXCL10/IP-10 and CCL5/RANTES were measured using duoset ELISA kits (R&D Systems), according to manufacturer’s instructions. Protein levels of all antimicrobial peptides were assayed using commercial ELISA assay kits, as previously described^12,22^. The sources of the individual ELISAs were as follows: Elafin (R&D Systems), Mannose binding lectin (MBL)-2 (R&D Systems) and SLPI (R&D Systems).

### RNA extraction, cDNA synthesis and quantitative PCR

Total RNA was extracted from sputum cell pellets or bronchial epithelial cell lysates stored in RLT buffer (RNeasy kit, Qiagen) and 2μg was used for cDNA synthesis (Omniscript RT kit). Quantitative PCR was carried out using previously described specific primers and probes for each gene of interest^23^, and normalized to 18S rRNA. Reactions were analysed using ABI 7500 Fast Realtime PCR system (Applied Biosystems, Carlsbad, CA).

### Virus detection, DNA extraction and bacterial 16S quantitative PCR

DNA extraction from total sputum was performed using the FastDNA Spin Kit for Soil (MP Biomedicals, Santa Ana, USA), as per manufacturer instructions. Bead-beating was performed for two cycles of 30 seconds at 6800 rpm (Precellys, Bertin Technologies, Montigny-le-Bretonneux, France). Total 16S bacterial loads were measured using quantitative PCR, as previously described^24^. Viruses were detected as described previously^12^.

### Statistical Analyses

Comparisons of sputum mRNA expression and protein concentrations between frequent and infrequent exacerbators were analysed using the Mann-Whitney U test. For *in vitro* experiments, baseline versus RV-induced expression was analysed using Wilcoxon rank-sum test. Mann Whitney *U* test was used to compare RV induction of mRNA or proteins between frequent and infrequent exacerbators. Correlations between datasets were examined using Spearman’s rank correlation coefficient. All statistics were performed using GraphPad Prism 6 software. Differences were considered significant when *P*<0.05.

## RESULTS

### Study population

We utilised a community-based cohort of COPD patients to evaluate anti-microbial immunity at baseline (stable-state) and during virus associated exacerbation. For baseline analyses of sputum cell mRNA (interferons (IFNs) and interferon-stimualated genes (ISGs)), 36 patients had sufficient sample for evaluation and, of these, 13 patients (36.1%) were classified as frequent exacerbators (clinical characteristics are shown in table 1). For baseline analyses of sputum soluble mediators (CXCL10/IP10, MBL, SLPI and elafin proteins), samples from a total of 50 COPD patients (combined from the community cohort and from subjects recruited to experimental infection studies) were available, with 15 patients (30%) classified as frequent exacerbators (clinical characteristics are also shown in table 1).

**Table 1:** Demographic and clinical characteristics of frequent and infrequent exacerbators included in baseline mRNA analyses and sputum soluble mediator analyses. GOLD = Global Initiative for obstructive lung disease

| Patients included in baseline mRNA analyses (n=36) |  |  |  |
| --- | --- | --- | --- |
|  | Frequent exacerbators (n= 13) | Infrequent exacerbators (n=23) | P value |
| Male sex | 10 (66.7%) | 27 (77.1%) | 0.49 |
| Age | 68.5 (58.3-75.3) | 67 (59.0-71.5) | 0.69 |
| GOLD stage I/II/III/IV | 0/9/3/3 | 5/28/2/0 |  |
| Long term oxygen use | 1 (6.7%) | 0 (0%) | 0.30 |
| Cardiac disease | 4 (26.7%) | 2 (5.7%) | 0.058 |
| Cerebrovascular disease | 3 (20%) | 1 (2.9%) | 0.075 |
| Diabetes | 2 (13.3%) | 2 (5.7%) | 0.57 |
| Pulmonary hypertension | 0 (0%) | 1 (2.9%) | 1.0 |
| Inhaled corticosteroid use | 9 (56.3%) | 11 (31.4%) | 0.11 |
| Beta agonist use | 14 (93.3%) | 18 (51.4%) | 0.0046 |
| Anti-muscarinic inhaler use | 12 (80.0%) | 12 (34.3%) | 0.0049 |
| Current smoker | 6 (40.0%) | 7 (20.0%) | 0.17 |
| Patients included in baseline sputum soluble mediator analyses (n=50) |  |  |  |
|  | Frequent exacerbators (n= 15) | Infrequent exacerbators (n=35) | P value |
| Male sex | 10 (66.7%) | 27 (77.1%) | 0.49 |
| Age | 68.5 (58.3-75.3) | 67 (59.0-71.5) | 0.69 |
| GOLD stage I/II/III/IV | 0/9/3/3 | 5/28/2/0 |  |
| Long term oxygen use | 1 (6.7%) | 0 (0%) | 0.30 |
| Cardiac disease | 4 (26.7%) | 2 (5.7%) | 0.058 |
| Cerebrovascular disease | 3 (20%) | 1 (2.9%) | 0.075 |
| Diabetes | 2 (13.3%) | 2 (5.7%) | 0.57 |
| Pulmonary hypertension | 0 (0%) | 1 (2.9%) | 1.0 |
| Inhaled corticosteroid use | 9 (56.3%) | 11 (31.4%) | 0.11 |
| Beta agonist use | 14 (93.3%) | 18 (51.4%) | 0.0046 |
| Anti-muscarinic inhaler use | 12 (80.0%) | 12 (34.3%) | 0.0049 |
| Current smoker | 6 (40.0%) | 7 (20.0%) | 0.17 |

### Evaluation of baseline expression of anti-viral mediators

Since viruses are a major cause of exacerbations and deficient anti-viral immunity has been observed in some but not all studies of COPD^8–11^, we hypothesised that frequent exacerbators would have reduced baseline expression of anti-viral immune mediators and thus an impaired potential to mount protective responses to virus infections. We examined baseline sputum cell mRNA expression of type I and III IFNs and ISGs in 36 patients from the community-based COPD cohort and found that frequent exacerbators had significantly reduced sputum *IFNβ* and *IFNλ2/3* mRNA expression with no difference in *IFNλ1* (Figure 1a). Baseline sputum concentrations of the interferon-inducible protein CXCL10/IP-10 (Figure 1b) were also reduced in frequent exacerbators but no difference in mRNA expression of other ISGs *2’-5’ OAS, viperin* or *M×1* was observed (Figure 1c).

**Figure 1:**
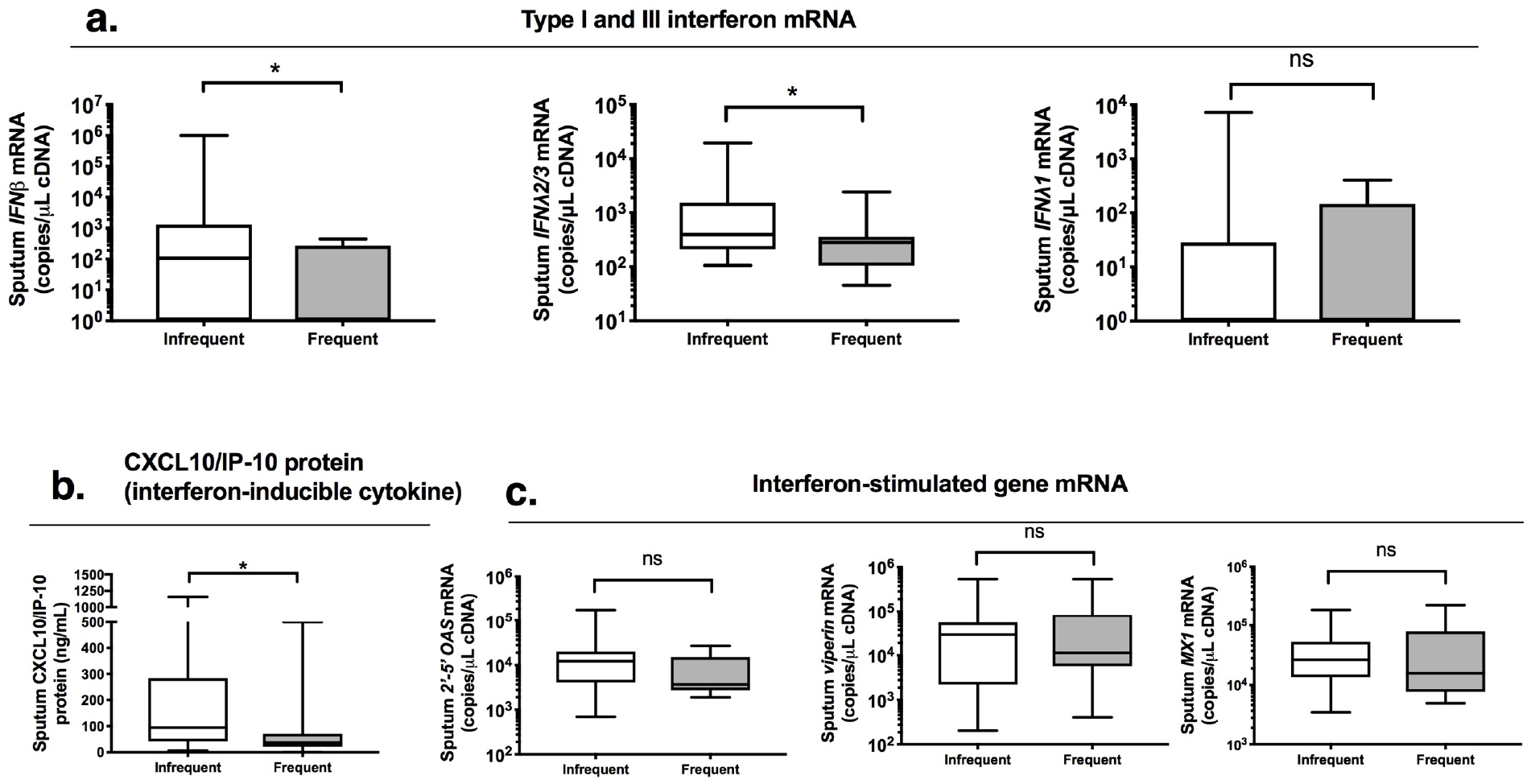
COPD patients with frequent exacerbations have reduced anti-viral immunity at clinical stability. A cohort of patients with COPD were monitored prospectively and sputum samples were taken when clinically stable for at least 6 weeks. Patients were stratified according to exacerbation frequency in the preceding year with patients who experienced ≥ 2 exacerbation episodes classified as ‘frequent’. Sputum cell mRNA expression of (a) *IFNβ, IFNλ2/3* and *IFNλ1* was measured by quantitative PCR. (b) Sputum supernatant protein concentrations of CXCL10/IP-10 were measured by ELISA. Sputum cell mRNA expression of interferon-stimulated genes (ISGs) (c) *2’-5’OAS, viperin* and *Mx1* was measured by quantitative PCR. Box and whisker plots showing median (line within box), IQR (box) and minimum to maximum (whiskers). Statistical comparisons were made using Mann Whitney U test. *p<0.05. ns = non-significant.

### Sputum interferon and ISG expression during virus positive exacerbation

In the community-based cohort, 18 episodes of acute exacerbation with positive virus detection were observed (rhinovirus n=11, coronavirus n=4, RSV n=2, influenza n=1). Of these, n=7 (38.9%) occurred in patients classified as frequent exacerbators. Clinical characteristics of the exacerbating patients are shown in table 2. One patient was hospitalised during exacerbation with all other episodes treated in the community. Having observed that frequent exacerbators have reduced capacity to mount an innate anti-viral response at stable state, we next examined sputum cell mRNA expression of IFNs and ISGs in the sub-group of patients who developed an exacerbation associated with positive virus detection. Frequent exacerbators had significantly reduced sputum cell expression of *IFNβ, IFNλ2/3 and IFNλ1* mRNAs at exacerbation presentation (Figure 2a). Sputum *2’-5’OAS* mRNA expression was also significantly reduced at exacerbation presentation in frequent versus infrequent exacerbators with no differences observed for *viperin* or *Mx1* expression (Figure 2b). Frequent exacerbators also had reduced sputum concentrations of the interferon-inducible cytokine CXCL10/IP-10 at exacerbation onset and 2 weeks post-onset with a similar non-significant trend (P=0.25) observed at exacerbation onset for CCL5/RANTES (Figure 2c).

**Figure 2:**
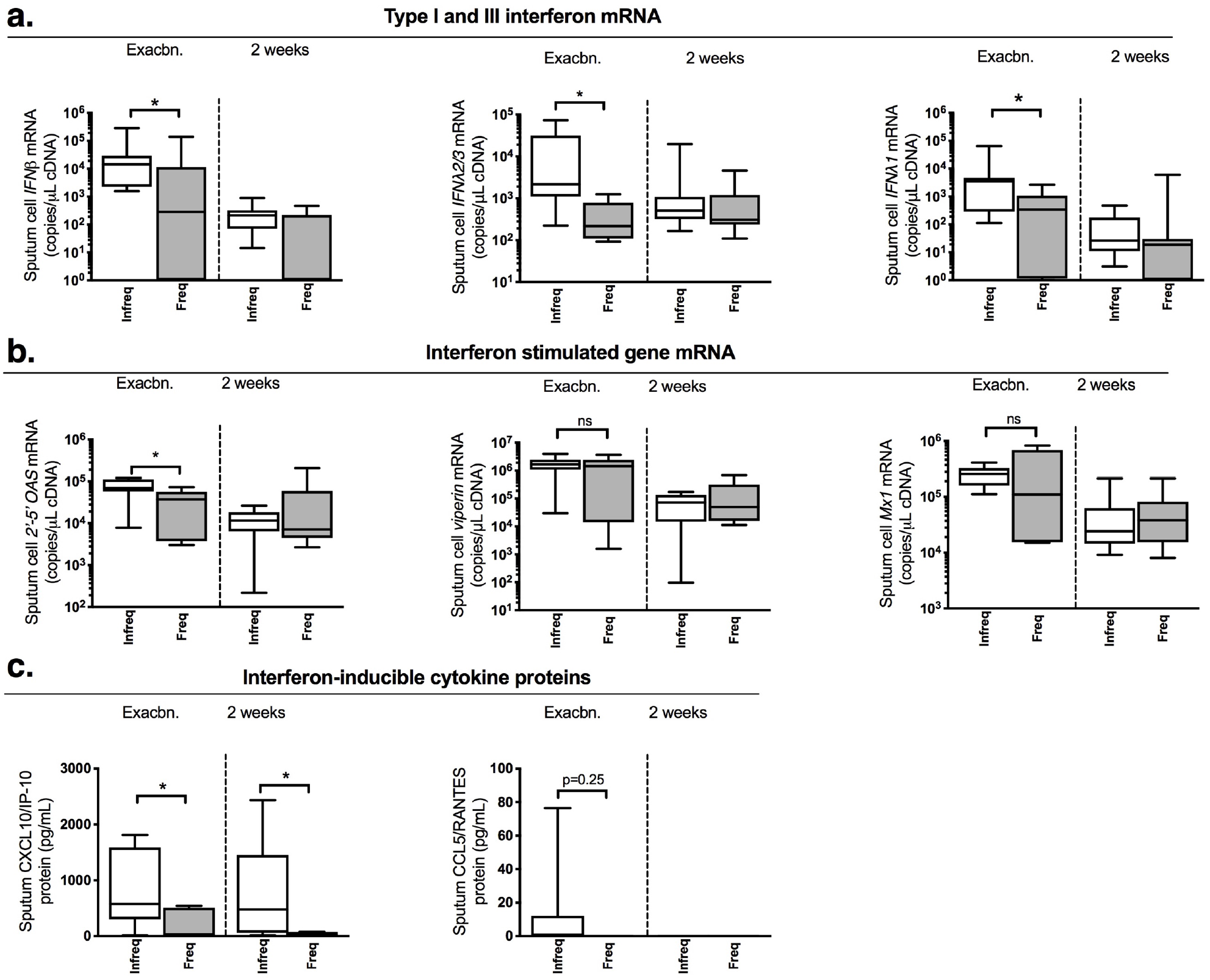
COPD patients with frequent exacerbations have reduced anti-viral immunity during virus-associated naturally occurring exacerbations. A cohort of patients with COPD were monitored prospectively. Patients were stratified according to exacerbation frequency in the preceding year with patients who experienced ≥ 2 exacerbation episodes classified as ‘frequent’. Sputum samples were taken at presentation with exacerbation associated with positive virus detection and 2 weeks during exacerbation. Sputum mRNA expression of (a) *IFNβ, IFNλ2/3* and *IFNλ1* (b) *2’-5’OAS, viperin* and *Mx1* was measured by quantitative PCR. Sputum protein concentrations of (c) CXCL10/IP-10 and CCL5/RANTES were measured by ELISA. Data displayed as box and whisker plots showing median (line within box), IQR (box) and minimum to maximum (whiskers). Statistical comparisons were made using Mann Whitney U test. *p<0.05. ns = non-significant.

**Table 2:** Characteristics of patients who developed virus associated exacerbation stratified according to exacerbation frequency.

|  | **Frequent exacerbators**<br>**(n=7)** | **Infrequent**<br>**exacerbations (n=11)** | **P value** |
| --- | --- | --- | --- |
| **Age**<br>**(median (IQR))** | 69 (64-74) | 71 (52-76) | 0.84 |
| **Male sex** | 3 (42.9%) | 9 (81.8%) | 0.14 |
| **GOLD Stage**<br>**(median (IQR))** | 2 (2-3) | 2 (1-2) | 0.13 |
| **Long term oxygen** | 0 (0%) | 0 (0%) | 1.0 |
| **Cardiac disease** | 0 (0%) | 1 (9.1%) | 1.0 |
| **Cerebrovascular**<br>**disease** | 1 (14.3%) | 0 (0%) | 0.39 |
| **Diabetes** | 1 (14.3%) | 1 (9.1%) | 1.0 |
| **Pulmonary**<br>**hypertension** | 0 (0%) | 0 (0%) | 1.0 |
| **Inhaled**<br>**corticosteroid use** | 4 (57.1%) | 3 (27.3%) | 0.33 |
| **Beta agonist use** | 6 (85.7%) | 7 (63.6%) | 0.60 |
| **Anti-muscarinic**<br>**inhaler use** | 4 (57.1%) | 6 (54.5%) | 1.0 |
| **Current smoker** | 4 (57.1%) | 2 (18.2%) | 0.14 |
| **Treatments during**<br>**exacerbation** |  |  |  |
| Antibiotics | 5 (71.1%) | 3 (27.3%) | 0.15 |
| Oral corticosteroids | 3 (42.9%) | 0 (0%) | 0.043 |

### *Ex vivo* bronchial epithelial anti-viral responses to RV infection are attenuated in frequent exacerbators

Having observed reduced IFN- and ISG-expression in frequent exacerbators from *in vivo* sputum samples taken during exacerbation, we next proceeded to determine whether epithelial cells from patients with COPD who are frequent exacerbators have deficient innate responses to RV infection. We utilised BECs collected bronchoscopically from sixteen patients with COPD (n=9 frequent exacerbators, n=7 infrequent exacerbators), differentiated at an air liquid interface and infected with RV-A1 at MOI = 0.1. Demographic/clinical characteristics of the included patients are shown in Table 3.

**Table 3:** Baseline characteristics of patients with COPD undergoing bronchoscopic sampling for airway epithelial cell experiments shown in figure 5.

|  | **Frequent exacerbators (n=9)** | **Infrequent exacerbations (n=7)** | **P value** |
| --- | --- | --- | --- |
| **Age** | 77 (72.5-82) | 67 (63.5 – 71) | 0.29 |
| **GOLD stage** |  |  |  |
| 2 | 1 (11.1%) | 2 (28.6%) | 0.55 |
| 3 | 8 (88.9%) | 6 (85.7%) | 1.0 |
| **Male sex** | 6 (66.7%) | 3 (42.9%) | 0.61 |
| **Inhaled corticosteroid use** | 7 (77.8%) | 3 (42.9%) | 0.30 |
| **Long acting Beta agonist use** | 7 (77.8%) | 3 (42.9%) | 0.30 |
| **Anti-muscarinic inhaler use** | 7 (77.8%) | 3 (42.9%) | 0.30 |
| **Current smoker** | 1 (11.1%) | 3 (42.9%) | 0.26 |

Greater heterogeneity with trend for increased viral load was observed in BECs from frequent exacerbators (Supplementary Fig 1). Significant induction of *IFNβ* and *IFNλ1* and *IFNλ2/3* mRNAs from baseline was observed in both frequent and infrequent exacerbators at 72 hours post-infection (Figure 3a). RV induction of *IFNλ1* mRNA was significantly lower in frequent versus infrequent exacerbators with a similar trend *(P=0.29)* observed for *IFNβ* mRNA but no significance difference observed for *IFNλ2/3* (Fig 3a). RV induction of the ISGs *2’-5’OAS* and *Viperin* mRNA was also significantly lower in frequent versus exacerbators with no significant difference observed for *PKR* (Figure 3b).

**Figure 3:**
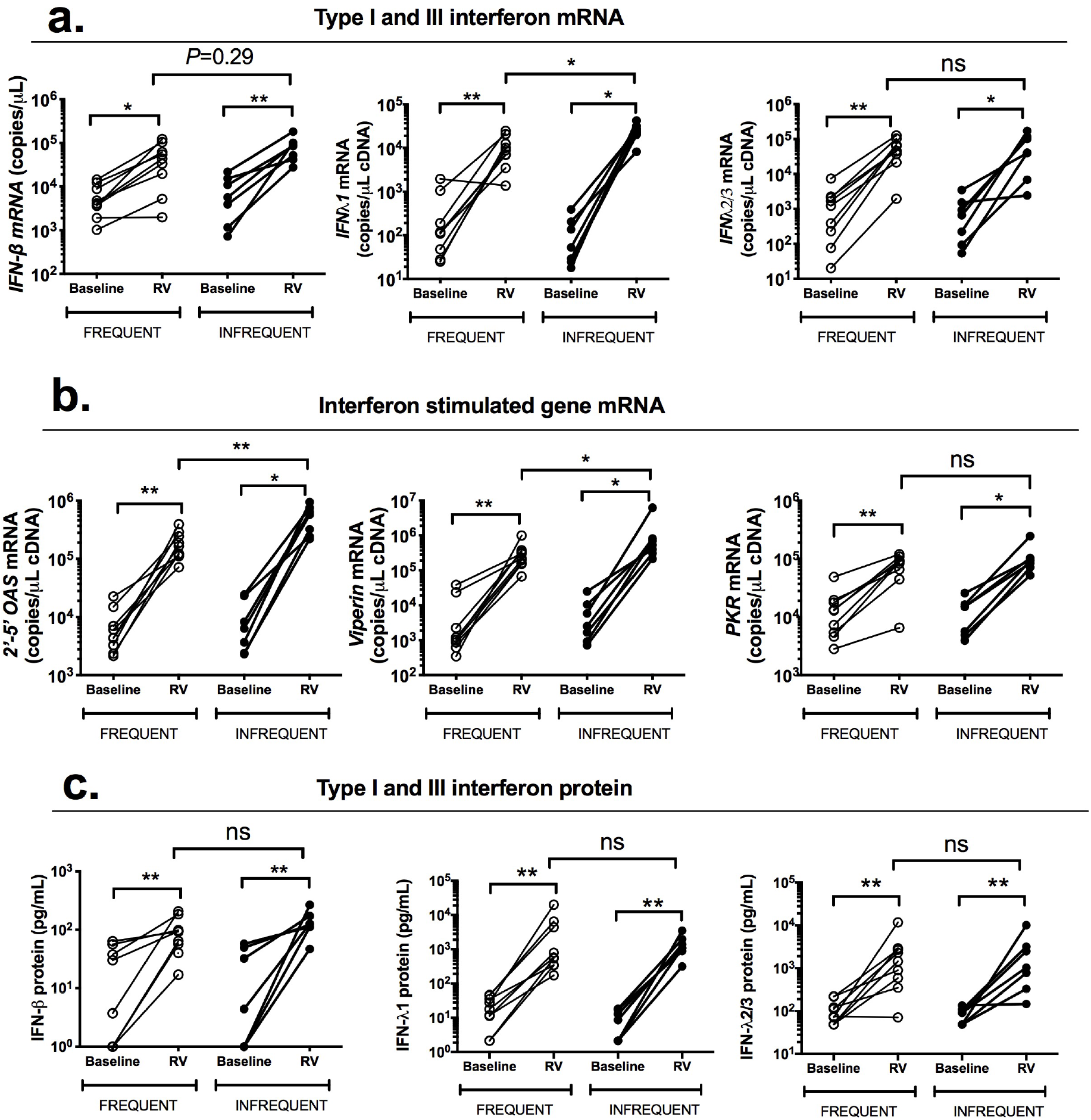
COPD patients with frequent exacerbations have deficient *ex vivo* innate anti-viral immune responses to rhinovirus infection in differentiated airway epithelial cells. Primary bronchial epithelial cells (BECs) from 16 patients with GOLD stage II or III COPD were differentiated at the air-liquid interface and infected *ex vivo* with rhinovirus (RV)-A1 or medium control (baseline). Cell lysates and supernatants were harvested post-infection. (a) *IFNβ, IFNλ1* and *IFNλ2/3 (b)2’-5’ OAS, PKR* and *viperin* mRNA expression in cell lysates at 72h was measured by quantitative PCR. (c) IFN-β, IFN-λ1 and IFNλ- 2/3 proteins were measured at 72h in cell supernatants by ELISA. Data represents individual patients and analysed by Wilcoxon rank sum test (baseline versus RV) or Mann Whitney *U* test (RV frequent exacerbators versus RV infrequent exacerbator). *p<.05; **p<0.01. ns = non-significant.

Significant induction of IFN-β, IFN-λ1 or IFN-λ2/3 proteins from baseline was also observed in both frequent and infrequent exacerbators at 72 hours post-infection (Figure 3c). However, there were no significant differences in the induction of IFN-β, IFN-λ1 or IFN-λ2/3 proteins by RV between frequent and infrequent exacerbators at 72 hours post-infection (Figure 3c).

Collectively, these data supported our *in vivo* findings from exacerbating patients that anti-viral responses are reduced in frequent exacerbators and extended these observations by identifying a dysregulated innate immune response in BECs defined by impaired expression of IFN and ISGs

### Evaluation of sputum bacterial loads at baseline and during virus-associated exacerbation

Virus-induced exacerbation can trigger subsequent bacterial infection in COPD^12,13^, a secondary phenomenon that is associated with increased exacerbation severity^12^ and inversely related to the interferon responses^21^. Having observed that anti-viral immunity is reduced in frequent exacerbators both *in vivo* and primary BECs from patients with COPD *in vitro,* we next examined whether sputum bacterial loads at stable state and during exacerbation differed according to exacerbation frequency. There were no differences in sputum 16S bacterial loads between frequent and infrequent exacerbators at baseline or exacerbation onset (Fig 4a&b). However, frequent exacerbators had a significant increase (~1 log) in bacterial loads at 2 weeks post exacerbation onset (Fig 4c), suggesting that this COPD subgroup may have increased likelihood of secondary bacterial infection during virus-induced exacerbation.

**Figure 4:**
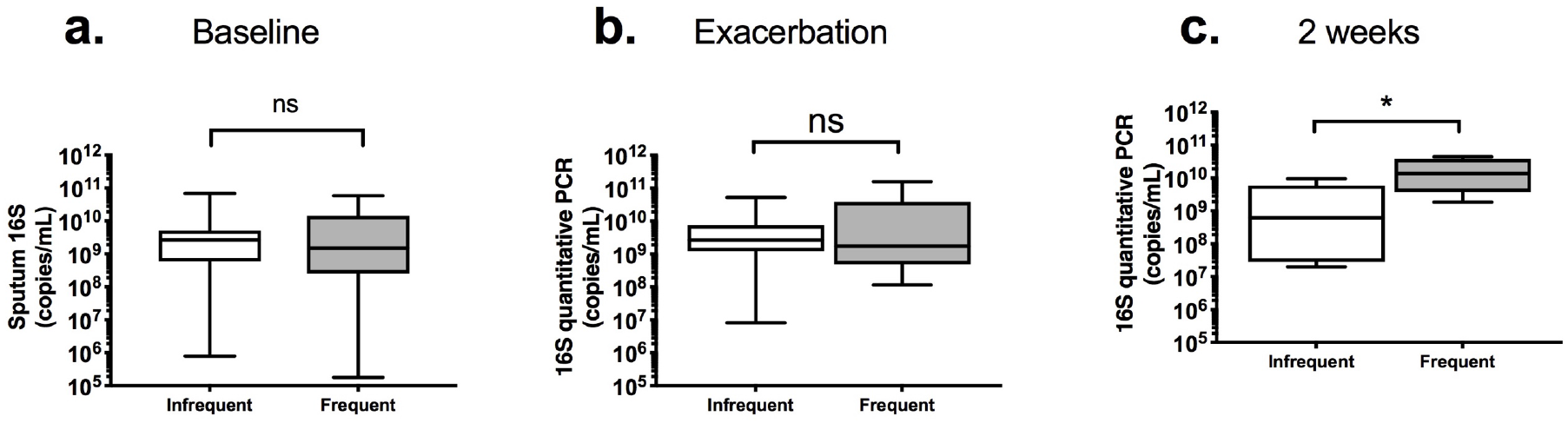
COPD patients with frequent exacerbations have increased bacterial loads during virus associated naturally occurring exacerbations. A cohort of patients with COPD were monitored prospectively. Patients were stratified according to exacerbation frequency in the preceding year with patients who experienced ≥ 2 exacerbation episodes classified as ‘frequent’. Sputum samples were taken at presentation with exacerbation associated with positive virus detection and 2 weeks during exacerbation. Sputum bacterial loads were measured by 16S quantitative PCR at (a) baseline (b) exacerbation and (c) 2 weeks. Data shown as median (IQR) and analysed by Mann Whitney U test. *p<0.05. ns = non-significant.

### No difference SLPI or elafin levels between frequent and infrequent exacerbators

We have previously reported that rhinovirus-induced secondary bacterial infection occurs through neutrophil elastase-mediated cleavage and reduction of the anti-microbial peptides (AMPs) SLPI and elafin, a process that occurs in COPD but not healthy subjects and may be further accentuated by inhaled corticosteroid use^21^. Given our observation of increased bacterial loads in frequent exacerbators, we next examined whether this sub-group of COPD patients have concurrently reduced SLPI and elafin levels. There were no differences in sputum SLPI or elafin levels between frequent and infrequent exacerbators either at baseline (Supplementary Fig 2a-b) or during exacerbation (Supplementary Fig 1c-d). This suggested that differential expression of these AMPs does not underlie increased secondary bacterial infection in frequent exacerbators.

### Airway mannose-binding lectin 2 levels are reduced in frequent exacerbators at baseline and during virus-associated exacerbation

We next focused on other another AMP that could be important in driving secondary bacterial infection in COPD. A previous study reported that mannose-binding lectin (MBL)-2 polymorphisms with associated MBL-2 deficiency in serum is associated with recurrence of infective exacerbations in COPD^25^. Consistent with this report, we also found that MBL-2 concentrations in sputum were significantly reduced in frequent versus infrequent exacerbators at stable state (Figure 5a) and also at 2 weeks post exacerbation onset (Figure 5b), the same timepoint at which bacterial loads were increased.

**Figure 5:**
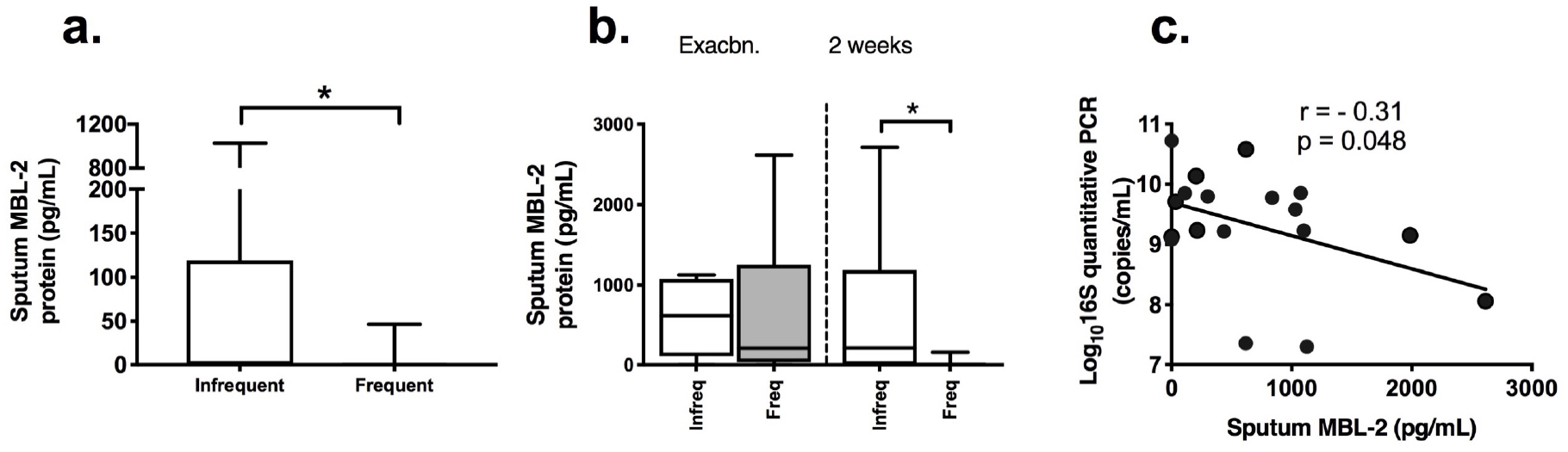
COPD patients with frequent exacerbations have reduced airway mannose-binding lectin 2 levels which correlate negatively with bacterial loads during virus associated exacerbations. Sputum protein concentrations of mannose binding lectin at (a) stable state (b) during virus associated exacerbation. (c) Correlation of sputum bacterial loads with sputum protein levels of mannose-binding lectin (MBL)-2. In (a)&(b) data displayed as box and whisker plots showing median (line within box), IQR (box) and minimum to maximum (whisker) and statistical comparisons were made using Mann Whitney *U* test. In (c), correlation analysis used was nonparametric (Spearman’s correlation). *p<0.05. ns = non-significant.

### Airway mannose binding lectin 2 levels negatively correlates with bacterial loads during virus-associated exacerbation

Having observed that frequent exacerbators have reduced MBL-2 levels and increased bacterial loads at 2 weeks post exacerbation onset, we next examined the relationship between these variables. There was a significant negative correlation between sputum concentrations of MBL-2 and bacterial loads (Figure 5c) suggesting that lower MBL-2 levels may contribute to increased secondary bacterial infection in frequent exacerbators.

## DISCUSSION

The underlying mechanisms involved in increased susceptibility to frequent exacerbations remains a crucial research question in COPD. We show here, for the first time, that patients with COPD who have a history of frequent exacerbations have reduced airway anti-microbial immunity when assessed at clinical stability and during subsequent virus-associated exacerbation with an associated increase in bacterial loads. We additionally confirm that COPD frequent exacerbators have reduced *ex vivo* immune responses to RV infection in primary airway epithelial cells, indicating an innate anti-viral deficiency. This is the first study to show that frequent exacerbators have impaired airway antimicrobial responses and provides an important mechanistic advance in our understanding of this clinically important COPD sub-group. These abnormalities could predispose such patients to greater risk of acquisition of pathogenic viruses and bacteria and/or promote a greater likelihood of developing an exacerbation following infection.

Previous studies have shown that *ex vivo* type I IFN responses to RV infection in bronchoalveolar cells and to influenza infection in BECs are impaired in COPD^8,9^. Patients with COPD also have increased virus loads and enhanced exacerbation severity when experimentally infected with rhinovirus^8,18^. However, defects in anti-viral immunity have not been shown universally with other studies reporting the opposite effect of augmented *ex vivo* anti-viral responses in COPD^10,11^. The discrepancy between these COPD studies suggests that defective immunity may not be present in all patients with a number of confounding competing factors such as disease severity and medications likely to have an influence. It is plausible that patients with frequent exacerbations might represent one sub-group with impaired anti-viral immunity. We report here that expression of both type I and type III IFNs, *IFNβ* and *IFNλ2/3* mRNAs is reduced in frequent exacerbators at stable state, suggesting that these patients might have a reduced potential to generate protective responses to virus infection. In support of this hypothesis, we also observed that anti-viral responses including *IFNβ, IFNλ1* and *IFNλ2/3* and the ISGs *2’-5’OAS* and CXCL10/IP-10 were suppressed during exacerbations in frequent exacerbators and that RV-induction of *IFNλ1, 2’-5’OAS* and *Viperin* mRNAs in BECs *ex vivo* was also reduced in frequent exacerbators.

Our study provides the first evidence that frequent exacerbators have innate anti-viral immune dysfunction, suggesting that a reduction in interferon production might underlie an augmented propensity to virus infections and thus increased exacerbation frequency observed in these patients. This mechanism is supported by a clinical study which reported that COPD patients who are frequent exacerbators experience significantly more coryzal episodes than infrequent exacerbators^26^. Given that inhaled IFN-β therapy has been shown to confer clinical benefit in a subgroup of severe asthmatic patients who develop a cold^27^, our data raises suggestion that such innate immune-boosting therapies could be effective when used in a targeted manner in patients with COPD with evidence of frequent exacerbations and a less effective innate immune response.

Despite observing impaired *ex vivo* RV-induction of IFN and ISG mRNA in frequent exacerbator BECs at 72 hours post-infection, we did not observe corresponding differences in protein production of type I and III IFNs by these cells at the same timepoint. Our studies were limited by sample availability only allowing evaluation at a single timepoint. We have previously reported that protein production of IFN persists at least up to 96 hours in this cell type^21^ and it is feasible that evaluation at later timepoints may be required to observe significant differences in translated IFN proteins. Despite a lack of difference in absolute levels of IFN protein, the reduced ISG expression observed in frequent exacerbators supports a less effective IFN-induced response in these subjects

Bacterial infection is also associated with exacerbation in COPD with increased PCR-based bacterial detection at exacerbation versus stable state suggesting a causative role^28,29^. Additionally, virus-induced secondary bacterial infection has been reported in both experimental and naturally occurring exacerbations^12,13^. We have previously reported that experimental RV challenge in patients with COPD is associated with increased frequency of secondary bacterial infection compared to healthy subjects^12,30^ via a mechanism of neutrophil elastase-mediated cleavage of anti-microbial peptides SLPI and elafin^12^. Here, we extend these findings to reveal that frequent exacerbators have higher bacterial loads at 2 weeks following onset of virus-associated exacerbation, suggesting that this subgroup of COPD patients might be at greatest risk of developing secondary bacterial infection following initial virus infection. Although we found no difference in levels of SLPI and elafin according to exacerbation frequency, we did identify that expression of another AMP MBL-2 was reduced in frequent exacerbators at stable state and during virus positive exacerbation. The potential importance of MBL-2 during virus-associated exacerbation was further demonstrated by our finding that its expression correlated negatively with bacterial loads during exacerbation. MBL is a key component of innate immunity that activates the lectin complement pathway and is involved in the process of bacterial phagocytosis^31^. MBL deficiency is associated with susceptibility to respiratory tract infections and pneumonia in healthy individuals^32,33^ and to increased risk of sepsis in animal models^34^. MBL can bind to various pathogens of relevance to COPD exacerbations including both viruses and bacteria^35^. Genetic MBL deficiency has been previously shown to be associated with both increased^25,36^ and reduced^37^ exacerbation frequency in COPD. Although serum MBL concentrations have been shown to not differ between frequent and infrequent exacerbators^36^, a previous study reported reduced bronchoalveolar lavage MBL concentrations in COPD versus healthy subjects^38^. Our data extends these observations by demonstrating that deficient airway MBL-2 at baseline and during subsequent exacerbation might predispose frequent exacerbators to increased secondary bacterial infection. A previous study reported that the presence of rhinovirus is associated with increased *S.pneumoniae* colonization in children with MBL gene polymorphisms, further suggesting a role for MBL in virus-induced secondary bacterial infection^39^. Given that nebulised plasma-derived MBL has been shown to restore phagocytic function in smoke-exposed mice^40^, our data raise speculation that administration of MBL could additionally be considered as an effective preventative and/or therapeutic strategy for COPD exacerbations. Further studies are needed to evaluate this.

It is important to note that causation cannot be inferred from the results of our study. Although we found reduced anti-microbial responses in patients with a history of frequent exacerbations, we cannot reliable conclude that these abnormalities directly lead to inherent increased exacerbation risk *per se.* There are likely to be a number of additional factors that may be contributing to the observed differences and, notably, in stable state analyses, frequent exacerbators showed trends towards greater ICS use and less current smoking. These confounders could contribute directly or indirectly to the differences observed, as previously reported^21,41,42^. We therefore consider our study to be hypothesis generating and emphasise the importance of future larger *in vivo* studies to more carefully dissect the influence of various factors and definitely answer the question of whether deficient antimicrobial immunity underlies propensity to exacerbation frequency in COPD. Previous studies have also indicated that exacerbation risk in COPD is dynamic and may change from year to year^4,43^. A longer-term study with repeated airway sampling would be need to determine whether baseline antimicrobial responses are similarly dynamic in nature.

In conclusion, we show that patients with COPD and a history of frequent exacerbations have reduced anti-viral and anti-bacterial immunity associated with increased secondary bacterial infection. These immune defects may underlie the increased propensity to exacerbations observed in this sub-group and provides evidence to support therapeutic approaches that boost innate immunity in COPD

## ABBREVIATIONS

BECs: bronchial epithelial cells
ALI: air liquid interface
COPD: Chronic obstructive pulmonary disease
FEV1: Forced expiratory volume in 1 second
FVC: Forced vital capacity
GOLD: Global Initiative for Obstructive lung disease
IFN: Interferon
ISG: Interferon-stimulated gene
MBL: Mannose-binding lectin
PEFR: peak expiratory flow rate
RV: rhinovirus
SLPI: Secretory leucocyte protease inhibitor

## Acknowledgements

**Funding:** Wellcome Trust Clinical Research Training Fellowship (grant number WT096382AIA) Imperial College Healthcare Trust Biomedical Research Centre grant (P33132); Academy of Medical Sciences and Wellcome Trust Starter Grant; Medical Research Council Program Grant G0600879; British Medical Association H.C. Roscoe Fellowships; BLF/Severin Wunderman Family Foundation Lung Research Program Grant P00/2; Imperial College and NIHR BRC funding scheme; the NIHR Clinical Lecturer funding scheme

## Author contributions

*Conception and Design:* AS, SLL, JF, SLJ, NWB, PM

*Experimental work:* AS, SLL, MC, MT, LJF, EB, JG, PV, PSP, KSN, AR, JF, NWB, PM

*Writing of manuscript:* All authors

*Guarantor:* AS

