## Supplementary Fig 1 for "Anti-microbial immunity is impaired in COPD patients with frequent exacerbations"

SUPPLEMENTARY FIGURES

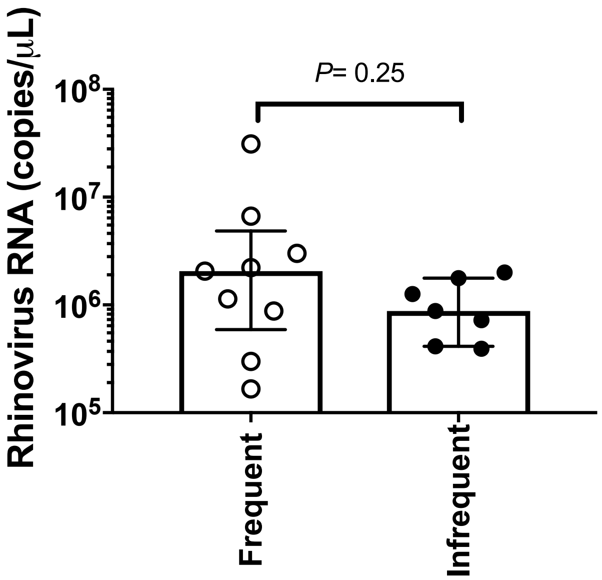


**Supplementary Figure 1: Virus loads following *ex vivo* infection of COPD bronchial epithelial cells in frequent- and infrequent-exacerbators.** Primary bronchial epithelial cells (BECs) from 16 patients with GOLD stage II or III COPD were differentiated at the air-liquid interface and infected *ex vivo* with rhinovirus (RV)-A1. Cells were harvested at 72 hours post-infection and rhinovirus RNA copies were measured by quantitative PCR. Data shown as median (IQR) and analysed by Mann-Whitney U test.

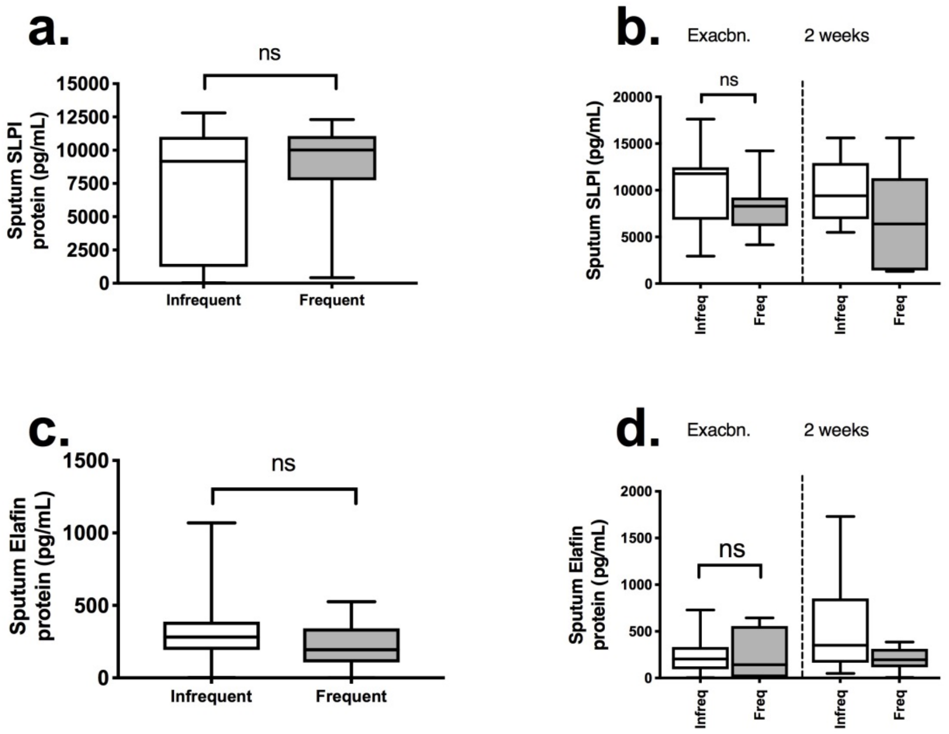


**Supplementary Figure 2: No difference in SLPI or elafin levels at clinical stability or during virus associated exacerbations between COPD frequent and infrequent exacerbators.** Sputum protein concentrations of secretory leucocyte protease inhibitor (SLPI) at (a) stable state and (b) during virus associated exacerbation were measured by ELISA. Sputum protein conentrations of elafin at (c) stable state and (d) during virus associated exacerbation were measured by ELISA. Data displayed as box and whisker plots showing median (line within box), IQR (box) and minimum to maximum (whisker) and statistical comparisons were made using Mann Whitney U test. ns = non-significant.
